# The effects of carnosine pretreatment on the inflammatory response and the PI3K/Akt signaling pathway following hypoxia-ischemia in neonatal rats

**DOI:** 10.1101/766063

**Authors:** Xiangmin Zhang, Lei Xia, Zhiheng Huang, Falin Xu

**Author notes:** Corresponding author (LX), (XMZ).

## Abstract

An increasing number of studies have demonstrated that carnosine plays a neuroprotective role in many types of brain injury. We have previously shown that carnosine has both short-term and long-lasting neuroprotective effects in a hypoxia–ischemia(HI) rat model. In the mature brain, post-ischemia neuronal survival involves in activation of the phosphatidylinositol 3-kinase (PI3K)/Akt signaling pathway, whether the activation of PI3K/Akt pathway also plays an important role in the immature brain still remain unclear.The goal of this study is to detect the effection of carnosine on inflammation response following HI, further evidencing neuroprotection of carnosine. We measured total Akt, phospho-Akt (p-AKT) and tumor necrosis factor receptor 1 (TNFR1) protein levels by western blot assay and tumor necrosis factor-α (TNF-α) and TNFR1 mRNA expression using real-time RT-PCR. We found the carnosine-pretreated group had statistically significant downregulation of TNF-α mRNA levels 24 h after HI (*P* < 0.05). Similar results were observed when we measured TNFR1 mRNA levels both 24h and 72h after HI (*P* < 0.05). And the TNFR1 protein expression after HI was markedly decreased at 24 and 72 h post-HI in the carnosine-pretreated rats(*P* < 0.05). Nevertheless, the rats pretreated with carnosine showed a marked increase in p-Akt levels (*P*< 0.05). And the pro-apoptotic protein Bad was also examined using immunohistochemistry after 24 and 72 h of all groups. We found significantly fewer Bad-positive cells in the carnosine-pretreated group at each time point after HI (*P* < 0.05). These findings suggest that carnosine pretreatment inhibits the HI-induced inflammatory response, and neuroprotection mechanism of carnosine involved in activation of the PI3K/Akt signaling pathway.

## Introduction

Perinatal hypoxic-ische infants mic(HI) brain injury is a main cause of neonatal mortality or permanent neurological dysfunction in surviving(1, 2). There is a growing body of evidence that post-ischemic inflammation plays a vital role in mediating the downstream effects of brain ischemia (3). Increased activity of tumor necrosis factor-α (TNF-α) is an important feature of brain injury and may be involved in the pathophysiology of such injury. The interaction of TNF-α with tumor necrosis factor receptor 1 (TNFR1) can induce apoptosis via a caspase-dependent pathway(4). Thus, we were interested in studying inhibition of inflammation as a potential strategy for alleviating neonatal HI-induced brain injury, thereby opening new avenues for clinical therapeutic intervention. Carnosine (β-alanyl-L-histidine) is a free radical scavenger present in a variety of organs, including the brain. Carnosine is well tolerated and safe (with no known side effects) and is commonly used as a dietary supplement(5, 6). It should be noted that many published studies have also provided compelling evidence that exogenously administered carnosine can exert a neuroprotective effect against brain injury resulting from a variety of causes(7, 8), this neuroprotection is believed to occur via the action of carnosine as a broad antioxidant(9). Moreover, the neuroprotective effect of carnosine pretreatment on HI-induced brain injury has been observed in a postnatal day 7 (P7) rat model (our unpublished data). However, most previously published experimental studies focused primarily on oxidative stress in the brain and neglected cerebral inflammation, which plays a central role in the mechanisms involved in brain injury following HI. Moreover, it is currently unknown whether the neuroprotection conferred by carnosine occurs via suppression of the inflammatory response and activation of the phosphoinositide 3-kinase/serine/threonine protein kinase (known collectively as PI3K/Akt) signaling pathway.

Akt is a critical survival factor that modulates cellular pathways involved in apoptosis. Akt inhibits apoptosis by phosphorylating and inhibiting pro-apoptotic mediators, including Bad(10). The PI3K/Akt signaling pathway is considered a central mediator in signal transduction pathways involving cell growth and cell survival, thereby maintaining the balance between cell survival and cell death(11). Traditionally, the neuroprotective function of PI3K/Akt has been attributed almost entirely to its anti-apoptotic actions(12). For example, PI3K/Akt-mediated neuroprotection in adult cerebral ischemia is well documented(13, 14). However, few studies have investigated whether the activation of the PI3K/Akt pathway plays an important role in the neuroprotection conferred by carnosine pretreatment following HI in the neonatal rat brain. Therefore, in this study, we investigated whether pretreatment with carnosine inhibits inflammation response and activate the PI3K/Akt signaling pathway in neonatal rat brain.

## Materials and Methods

### General preparation

Sprague–Dawley 7-day-old (P7, with P0 taken as the day of birth) rat pups (12–18 g) of both genders were used. The animals were purchased from Shanghai Experimental Animal Center of Chinese Academy of Sciences, and were randomly assigned to the following three groups containing 17 pups each, 1) sham control group (no carotid ligation and hypoxia), 2) HI-only group (carotid ligation and hypoxia), and 3) pretreatment group (carnosine pretreatment followed by carotid ligation and hypoxia). Pups in each group were chosen from among different litters to maintain parity within the groups. L-carnosine (Sigma, USA) was dissolved in normal saline. In the carnosine pretreatment group, carnosine was administered by intraperitoneal injection at a dosage of 250 mg/kg (injection volume of 0.25 ml/kg)(15) 30 min prior to the induction of HI. Members of the HI-only group were pretreated with an equal volume of normal saline. The sham control group did not receive any treatment.

### Hypoxic-ischemic (HI) model

All procedures were conducted in strict accordance with Directive 86/609/EEC on the protection of animals used for scientific purpose. The Animal Ethics Committee of Fudan University approved all procedures using animals. The HI model was induced as previously described(16), with slight modifications described below. Briefly, pups were housed with a dam under a 12h light/12h dark cycle with food pellets and tap water available *ad libitum* throughout the study. P7 rats were anesthetized with anhydrous diethyl ether. The left common carotid artery was exposed, isolated and cut between two sutures. After surgery, the pups recovered for 1.5 h. The pups were then placed in a humidified chamber partially submerged in a 37°C water bath. HI was induced by perfusing the chamber with humidified 8% oxygen balanced with 92% nitrogen for 2 h. After hypoxic exposure, the pups were returned to their biological dams. This protocol resulted in moderate brain injury. Sham-operated controls were randomly chosen from the same litters as the HI rats and underwent anesthesia and incision only. Throughout the study, all efforts were made to minimize the number of animals used and their suffering.

### Quantitative real-time PCR assays

To determine the mRNA levels of TNF-α and TNFR1, animals were killed by decapitation 24 h after HI (n = 4 per group). Total RNA was extracted using Trizol (Invitrogen, USA) according to the manufacturer’s instructions. RNA was quantified by ultraviolet absorption at 260 nm. cDNA was synthesized with the SuperScript Reverse Transcriptase kit (Promega, USA) according to the manufacturer’s instructions. Quantitative real-time PCR was performed in the Eppendorf Mastercycler ep realplex^4^ (Eppendorf, Germany) as described previously(16) using the following primers (based on the accession numbers indicated): TNF-α (NM-012675) forward: 5’-CTCATTCCTGCTCGTGGC-3’, reverse: 5’-TCTGAGTGTGAGGGTCTGGG-3’,TNFR1 (AF_329978) forward, 5’-GCAAGGACC CAAACAAAG-3’, reverse: 5’-TTCAACCGTTCATCCATTA-3’, and β-actin (NM_031144) forward: 5’-TGTCACCAACTGGGA CGATA-3’, reverse: 5’-CAGCATGTGTTGGCATAGAGGTC-3’. Each sample was processed three times in duplicate. After the final cycle, a melting curve was determined to verify that only one product had been produced. The amplified transcripts were quantified by the comparative C_T_ method using the standard curves with β-actin as an internal control.

### Western blot assays

For western blot analysis, brain tissues were homogenized in 0.7 ml of RIPA buffer containing 50 mM Tris (pH 7.4), 150 mM NaCl, 1% NP-40, 0.5% sodium deoxycholate, 0.1% sodium dodecyl sulfate, 0.5 mM phenylmethanesulfonyl fluoride (PMSF), sodium orthovanadate, sodium fluoride, ethylene diamine tetraacetic acid (EDTA) and leupeptin(n = 4 per group). The proteins were then transferred onto a nitrocellulose membrane (Millipore Corporation). After blocking by rotating in 5% bovine serum albumin (BSA) or 10% skim milk powder in TBS-T for 1h at room temperature, the membranes were incubated with the following primary antibodies (at the indicated dilutions) overnight at 4°C, anti-phospho-Akt (p-Akt) (Ser473, Cell Signaling Technology, 1:500), anti-Akt (Cell Signaling Technology, 1:200) and TNFR1 (Santa Cruz Biotechnology; 1:200). GADPH was used as an internal control for protein quantification (KangChen, Shanghai, China, 1:3000). After washing, the membranes were incubated with horseradish peroxidase-conjugated goat anti-rabbit IgG (Santa Cruz Biotechnology, 1:3000). Immunoreactive proteins were then visualized using the SuperSignal West Pico Trial ECL enhanced chemiluminescence detection kit (Pierce, USA) combined with a western blotting detection system (FUJIFILM, Japan).

### Tissue preparation and immunohistochemistry

24 or 72 h after HI induction, the pups were weighed and anesthetized with anhydrous diethyl ether (n = 5 per group). Coronal paraffin sections were cut approximately 3um from the bregma at the level of the hippocampus; the location was confirmed by hematoxylin and eosin staining. Briefly, the sections were dewaxed in dimethylbenzene and dehydrated through 100%–70% graded ethanol in distilled water. Endogenous peroxidase was inhibited with 0.3% hydrogen peroxide in methanol at room temperature for 10 min. After washing with PBS and digesting with trypsin for 15 min at 37°C, the sections were incubated in primary antibody (anti-Bad, Biovision, USA, 1:50) in blocking solution at 4°C overnight. After washing out the unbound primary antibody, the sections were incubated with goat anti-mouse IgG for 60 min at room temperature. Immunoreactivity was visualized with a mixture containing 0.05% 3, 3′-diaminobenzidine (DAB) and 0.03% H_2_O_2_ for 5 min. For each slice, three fields of view of the hippocampal CA1 region were photographed using a stereological fluorescence microscope (Leica, Germany). Cells with brown cytoplasmic coloring were considered to be positive for Bad. The number of Bad-positive cells in each field of view was counted under 400× magnification by an examiner who was blind with respect to the group assignment.

### Statistical methods

All quantitative data are presented as mean ± standard deviation (S.D.). The significance of differences between groups was determined by one-way analysis of variance (ANOVA) followed by Tukey’s HSD or Dennett’s T_3_ test (where equal variance was not assumed), and differences with a *p*-value less than 0.05 were considered statistically significant. All analyses were performed using SPSS 19.0 statistical software.

## Results

### Pretreatment with carnosine alleviated the HI-induced inflammatory response

To investigate whether carnosine pretreatment decreases the inflammatory response after HI, we measured hippocampal TNF-α and TNFR1 mRNA levels using real-time RT-PCR analysis. Relative to the HI-only control group, the carnosine-pretreated of HI group had statistically significant downregulation of TNF-α mRNA levels 24 h after (Fig 1, *F* = 100.7, *P* = 0.001). Compared with the sham operation group, the difference was not statistically significant(P>0.05). And after 72 h, it had returned to the control levels in the sham group(Fig 1, *F* = 1.51, *P* =0.272). At the same time, the level of TNFR1 mRNA in HI group was significantly higher than that of carnosine-pretreated HI group(Fig 2A, F = 381.19,P =0.001, 24 h; F = 2.40, P =0.176, 72h). Similar results were observed when we measured TNFR1 expression (Fig 2A, *P* < 0.05). We also examined TNFR1 protein levels using western blot analysis. Our findings showed that TNFR1 protein levels in HI-only control rats were higher than those in the sham control group 24 h after HI (Fig. 2B, *F* = 35.38, *P* = 0.001, 24h). Unlike the mRNA analysis, however, TNFR1 protein levels in the HI group remained elevated at 72 h (Fig 2B, *F* = 23.02, *P* = 0.001). Nevertheless, TNFR1 protein expression following HI was markedly decreased at 24 and 72 h post-HI in the carnosine-pretreated rats.

**Fig 1.**
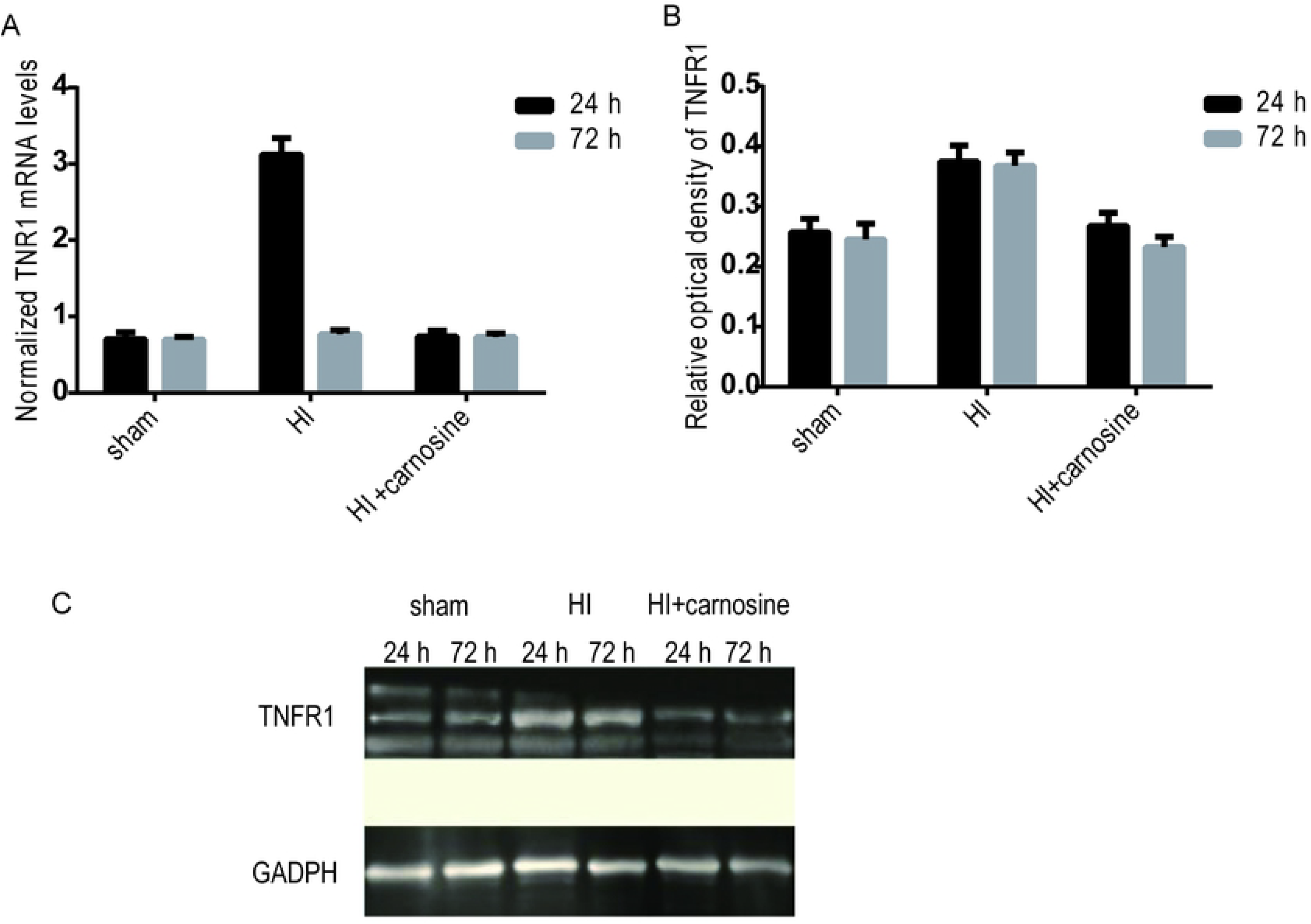
Carnosine pretreatment decreases hippocampal TNF-α mRNA levels in neonatal rats after HI. TNF-α mRNA was quantified by the comparative C_T_ method, employing β-actin as the internal control. * *p* < 0.05 vs. HI control group at 24 h.

**Fig 2.**
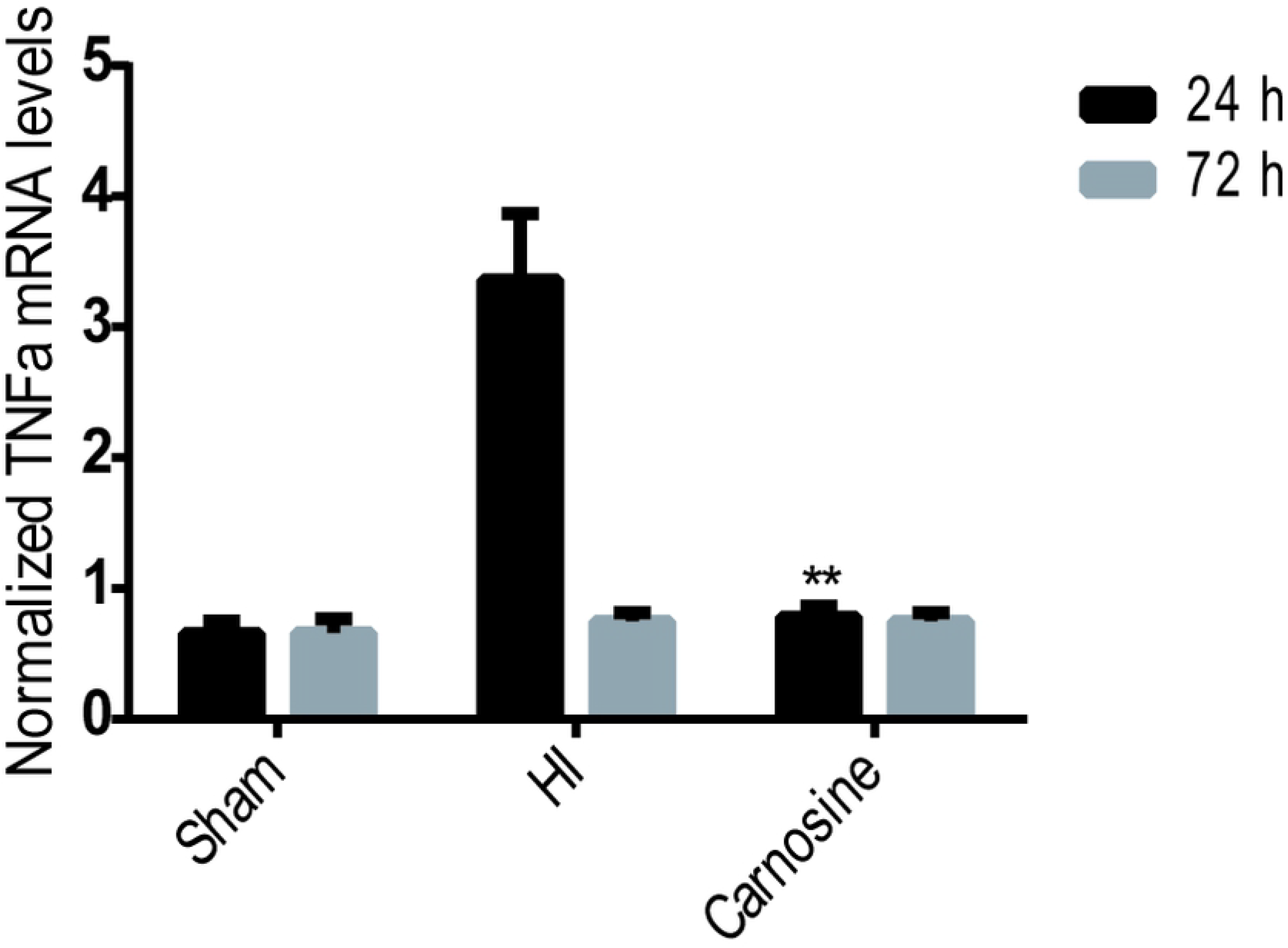
Carnosine pretreatment decreases hippocampal TNFR1 mRNA and protein levels in neonatal rats after HI. (A) At the mRNA level. * *p* < 0.05 vs. HI control at 24 h. (B) At protein levels. * *p* < 0.05 vs. HI control group at 24 and 72 h. (C) Western blot results. The samples of each group were separated by electrophoresis at different time points

### Pretreatment with carnosine protects the neonatal brain via activation of Akt

To determine whether carnosine pretreatment activates Akt in neonatal rats following HI, we used western blot analysis to measure p-Akt and total Akt protein levels in the hippocampus 24h and 72h after HI were induced. Compared with the HI-only control animals, rats pretreated with carnosine showed a marked increase in p-Akt levels (Fig 3A, *P* < 0.05 (*F* = 26.38, *P* = 0.001, 24 h; *F* = 22.62, *P* = 0.001, 72 h),). However, total Akt protein levels were not changed in each group at the different time points following HI (Fig 3B, *F* = 0.14, *P =* 0.84, 24 h; *F* = 0.05, *P =* 0.95, 72 h, *P* > 0.05). And there was no statistically significant difference between the pretreatment group and the sham operation group (*P* > 0.05).Furthermore, the total Akt protein level was compared at different time points in each group, and the difference was not statistically significant (*P* > 0.05).

**Fig 3.**
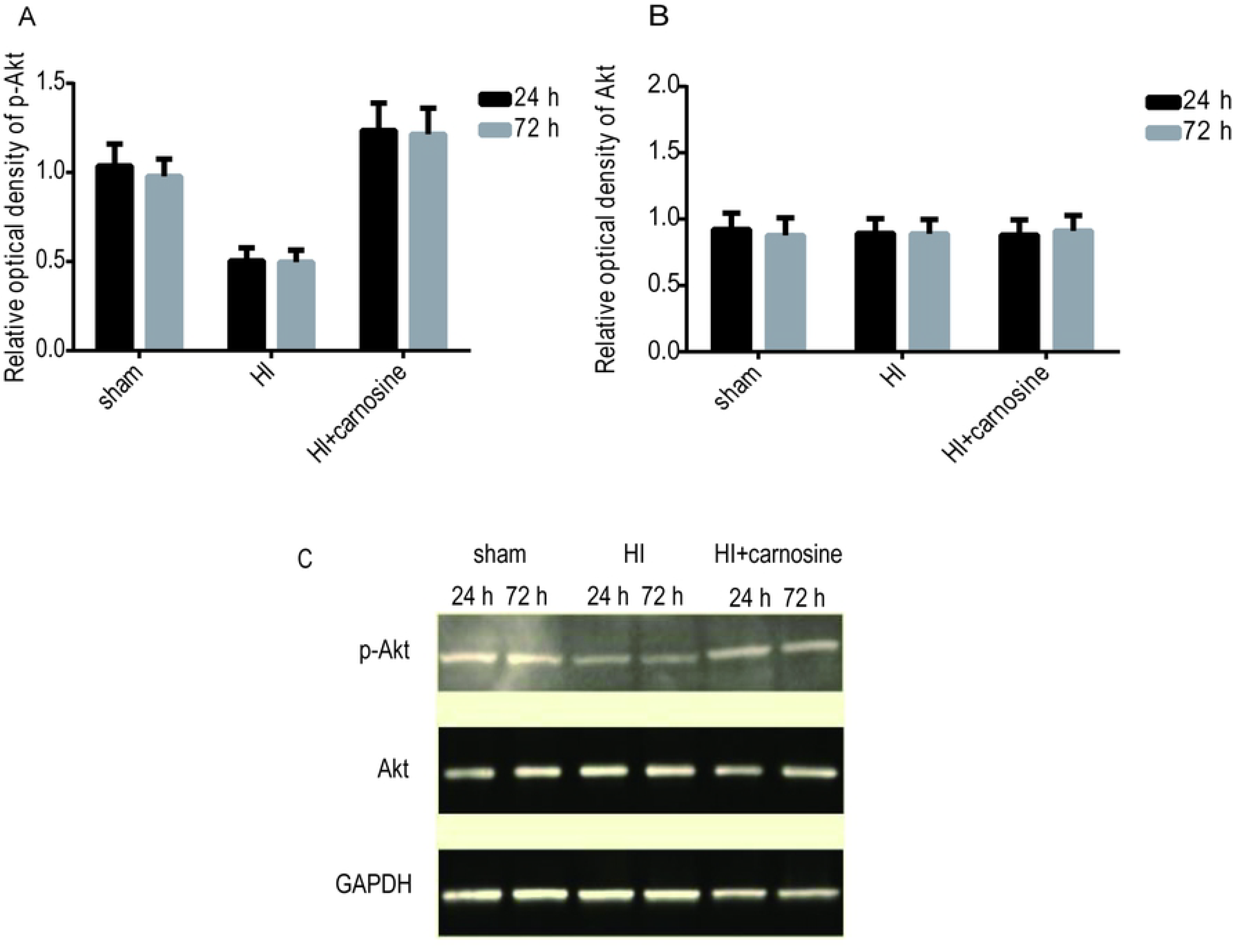
Carnosine pretreatment elevates p-Akt but not total Akt levels in P7 rats after HI. (A) Activated Akt (p-Akt) protein levels. * *p*<0.05 vs. HI-only control group at 24 and 72 h. (B) Total Akt protein levels. (C) Western blot results. The samples of each group were separated by electrophoresis at different time points

### Pretreatment with carnosine decreases expression of the pro-apoptotic protein marker Bad following HI

To test whether Akt exerts neuroprotective effects against apoptosis via inhibition of the pro-apoptotic protein Bad, Bad protein levels were examined by immunohistochemistry 24 and 72 h after HI (Fig 4). Consistent with our hypothesis, we found significantly fewer Bad-positive cells in the carnosine-pretreated group compared with the HI-only control group at each time point following HI (Fig 4, *P* < 0.05). Immunohistochemical results showed that the number of BAD positive cells in HI group without pretreatment of myotide was higher than that of HI group (Fig 4, F = 141.14, *P* = 0.001, 24h; F =153.31, *P* = 0.001, 72h).Carnosine different time points after pretreatment significantly reduce the HI BAD positive cells number, compared with HI group, the difference was statistically significant (Fig 4, *P* < 0.01), compared with between the control group, there was no statistically significant difference (*P* > 0.05).This is consistent with the different expression trends of p-akt in the pretreatment group and HI group.

**Fig 4.**
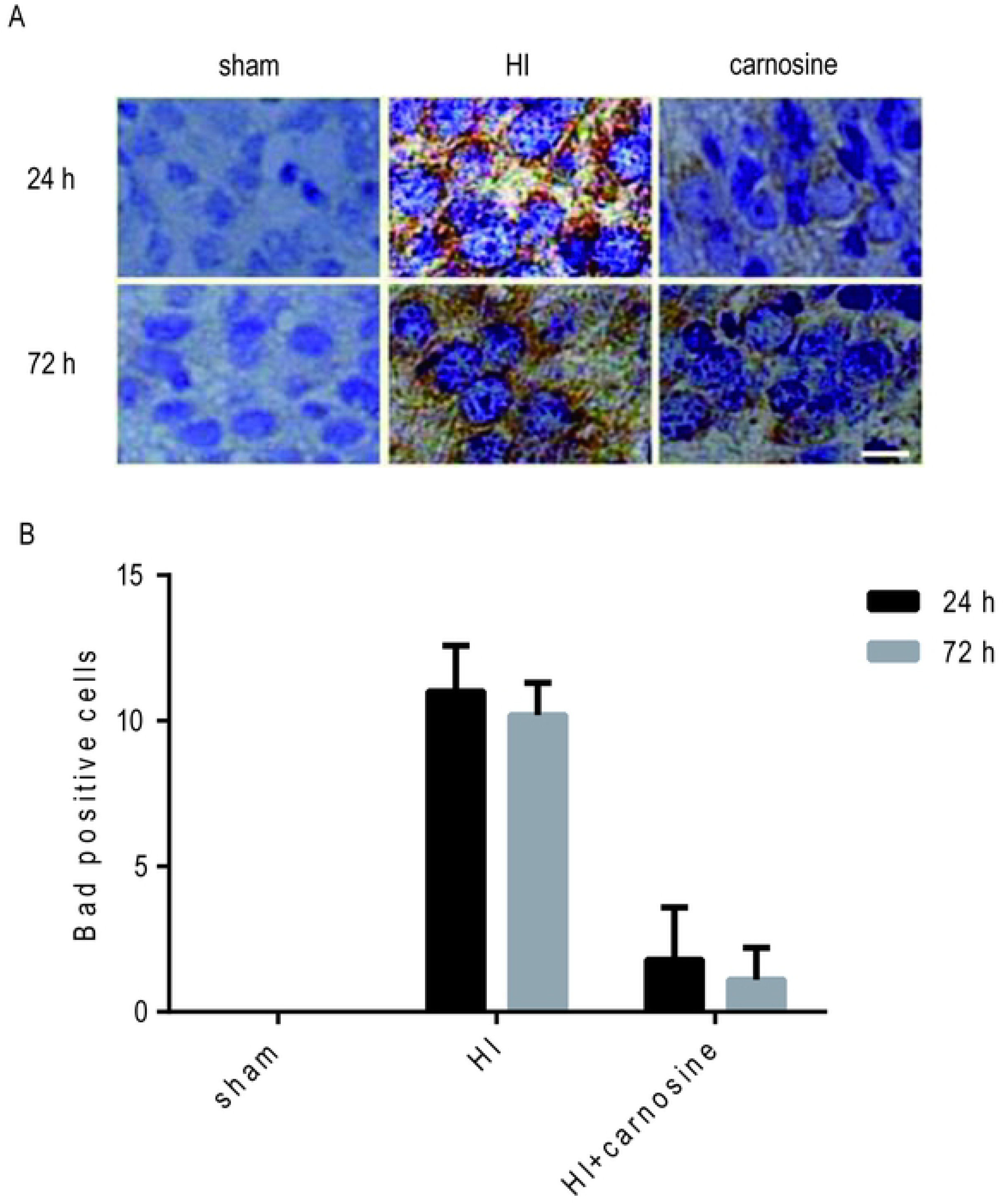
Carnosine pretreatment decreases the number of Bad-positive cells in neonatal rats after HI. (A) Color micrographs of the hippocampus at 400× magnification. (B) Bad-positive cells counts. * *p* < 0.05 vs. HI control group at 24 and 72 h.

## Discussion

In the present study we used a well-established P7 rat HI model to investigate the effects of carnosine on the inflammatory response and the PI3K/Akt signaling pathway. The obtained results demonstrated that the neuroprotective effect of carnosine pretreatment against HI-induced brain injury was mediated by both suppressed inflammation and activation of the PI3K/Akt signaling pathway.

Previous study has shown that secondary inflammation after HI has been shown to contributes to HI-induced brain injury(17). The toxic effects of inflammatory cytokines include both direct free radical release and indirect excitatory amino acid production(18).

TNF-α is a cytokine involved in different pro-inflammatory actions. Moreover, TNF-α expression is increased in a wide range of CNS disorders, including ischemia(19). An *in vitro* study has shown that exposing cells to TNF-α leads to apoptosis by activation of a caspase cascade(20). Most cytotoxic effects of TNF are mediated by the interaction of TNFR1 with the death domain (TRADD) protein of TNF death domain(21).Tumor necrosis factor death domain protein (TRADD) interacts with fas-related death domain protein (FADD) on caspase-8 and subsequently activates caspase-3, thus initiating the apoptosis pathway. The expression and release of TNF dramatically increase under pathological conditions. In some cases, the expression of TNF-appeared after brain injury and 1h before neuron death(22), while TNF is activated after binding to the TNFR1 receptor on neurons and glial cells of the central nervous system(23).The activation of TNFR1 receptor activates the caspase pathway and induces apoptosis(24).Therefore, TNF-overexpression has obvious cytotoxic effects. In our P7 rat model, TNF-α mRNA levels increased 24 h after HI and then declined to control levels by 72 h post-HI. Pretreatment with carnosine markedly decreased TNF-α mRNA levels after HI compared with untreated rats.

The mRNA levels of the TNFR1 were also examined 24 and 72 h after HI and obtained results were consistent with TNF-α mRNA levels, which is in line with TNF-α exerting its biological function via interaction with TNFR1. Interestingly, TNFR1 protein levels remained elevated even at 72 h post-HI, yet, the reasons causing the difference between mRNA and protein changes remain unclear. Nevertheless, the decreased TNF-α mRNA, TNFR1 mRNA and TNFR1 protein levels in the carnosine-pretreated group early after HI may have helped to alleviate HI-induced brain injury.

Akt has profound effects on the regulation of cell metabolism. Although the stimulation of Akt mainly alleviates the activation of the endogenous apoptotic pathway, carnosine can theoretically inhibit the inflammatory response due to its antioxidant effect.Previous studies have shown that activation of Akt signaling pathway has an important role in cerebral ischemia, hypoxic ischemia and ischemia reperfusion injury. Existing studies have shown that the infarction rate of transgenic mice with p-akt overexpression is reduced by 35% after ischemia reperfusion(25).Recently, it has been shown that cerebral ischemia for 10min followed by 10min reperfusion can reduce the cerebral infarction rate by 50%. It has also observed that the neuroprotective effect of reperfusion is mediated by activation of Akt signaling pathway(26).Our previous studies have also shown that carnosine preconditioning has a neuroprotective role by alleviating the apoptosis of nerve cells in P7 ischemic rats, reducing the rate of infarction and improving the long-term neurological results(27).

Our data suggest that the activation of Akt is critical for carnosine pretreatment–induced neuroprotection against HIBD. Akt confers neuroprotection by inactivating the pro-apoptotic Bcl-2 family member Bad through PI3K/Akt-mediated phosphorylation(28). Unphosphorylated Bad promotes apoptosis by binding and neutralizing anti-apoptotic Bcl-2 proteins. Many studies have confirmed that p-akt level decreases and total Akt levels remains unchanged in the case of ischemia(29).In this study, intraperitoneal injection of carnosine prior to HI induction had a beneficial effect. Carnosine preconditioning prevented hypoxic-ischemia-induced reduction of p-akt levels, while total protein levels remained unchanged in each group, which is consistent with previous reports. BAD is a key molecule that conveys information of the extracellular environment through phosphorylation(30).Non-phosphorylated BAD induces apoptosis by damaging the mitochondrial membrane, resulting in the release of cytochrome C from the mitochondrial membrane into the cytoplasm to activate the caspase-9 or apoptotic protease-activated factor-1 complex(31).

Our study provides evidence that activation of the Akt signaling pathway is critical to the neuroprotection of HI brain injury induced by carnosine preconditioning.Its mechanism of action is the BAD inactivation of bcl-2 family members via mediated phosphorylation, which has a neuroprotective role. This study showed that carnosine pretreatment reduced the number of BAD positive cells in the hippocampus at different time points after HI, which further confirmed that Akt promoted cell survival through phosphorylation of BAD pro-apoptotic protein.Although we only detected the expression of BAD protein by immunohistochemistry, we indirectly speculated the role of Akt signaling pathway in the BAD phosphorylation process.

In this study, administering carnosine prior to inducing HI conferred a beneficial effect, as carnosine pretreatment prevented HI-induced decreases in p-Akt levels. Interestingly, however, total Akt protein levels were unchanged. In addition, compared to HI-only rats, carnosine-pretreated rats had markedly fewer Bad-positive cells in the hippocampus at each time point following HI, which further demonstrated that Akt confers an anti-apoptotic effect by inactivating the pro-apoptotic protein Bad.

To sum up, our findings show the effect of carnosine on the inflammatory response in the developing rat brain following hypoxia-ischemia. To the best of our knowledge, this is the first study to demonstrate a relationship between carnosine intervention and inflammation and PI3K/Akt signaling. These findings strongly suggest that the neuroprotective effects of carnosine pretreatment are rooted in inhibition of inflammation after HI and are mediated by the PI3K/Akt signaling pathway.

## Acknowledgments

This study was supported in part by the Neonatal Disease Key Laboratory of the Ministry of Public Health from Fudan University in China. The authors are grateful to the faculty of the pediatric institute for excellent technical assistance.

## Author Contributions

Conceived and designed the experiments: XZ XC. Performed the experiments: XZ LX ZH. Analyzed the data: XZ LX ZH. Contributed reagents/materials/analysis tools: XZ XC FX. Wrote the paper: XZ LX.

